# Promising effects of duck vaccination against highly pathogenic avian influenza, France 2023-24

**DOI:** 10.1101/2024.08.28.609837

**Authors:** Claire Guinat, Lisa Fourtune, Sébastien Lambert, Eva Martin, Guillaume Gerbier, Andrea Jimenez Pellicer, Jean-Luc Guérin, Timothée Vergne

## Abstract

Highly pathogenic avian influenza causes significant poultry losses and zoonotic concerns globally. Our assessment of duck vaccination in France predicted that 314-756 outbreaks were averted in 2023-24, representing a 96-99% reduction in epizootic size, likely attributable to vaccination.

## Text

Highly pathogenic avian influenza (HPAI) H5 viruses of clade 2.3.4.4b continue to affect diverse regions and species worldwide. Since 2020, this ongoing HPAI panzootic has reached an unprecedented scale, leading to the death or culling of over 130 million poultry across 67 countries, substantially threatening food security^1^. Mass mortality events in wild birds and spillovers to over 48 mammal species across 26 countries have raised concerns about both conservation and zoonotic pandemic risks ^2^.

While most countries rely on poultry depopulation and movement restrictions to control HPAI, France has recently adopted a complementary preventive vaccination strategy^3^. Since October 2023, domestic ducks in the production stage are being vaccinated with the inactivated vaccine Volvac B.E.S.T. AI + ND (Boehringer Ingelheim), receiving two doses at 10 and 28 days, and a third dose at 56 days in high-risk zones and winter periods^4^. Since May 2024, the RESPONS AI H5 vaccine (Ceva Animal Health), a self-amplifying mRNA vaccine, was also added to the vaccination campaign. Vaccination for breeder duck flocks remains optional. As of July 1, 2024, more than 35 million ducks have received two vaccine doses and 1.5 million have received all three vaccine doses^4^.

In 2023-24, only 10 poultry farm outbreaks of HPAI H5 were reported, representing a substantial reduction from the 1,374 outbreaks detected in 2021-22 and 396 outbreaks in 2022-23 (**Figure 1A**). In contrast, outbreaks continued in non-vaccinating European countries (**Figure 1B** and **1C**). Despite these encouraging results, the extent to which vaccination contributed to this reduction remains unclear.

**Figure 1.**
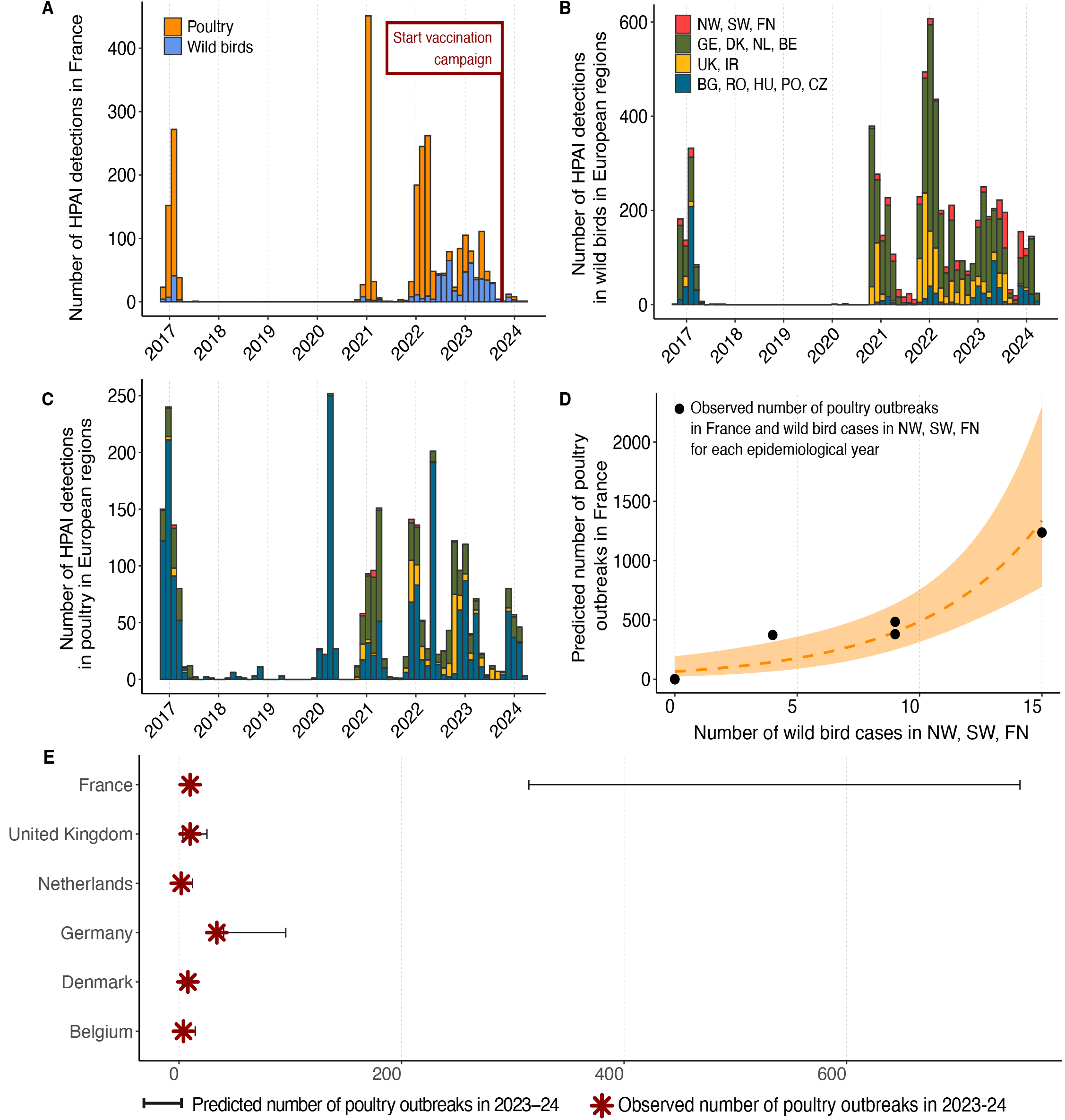
**A)** Temporal distribution of highly pathogenic avian influenza (HPAI) H5 (clade 2.3.4.4b) poultry farm outbreaks (orange) and wild bird cases (blue) in France. The start of the vaccination campaign (October 1^st^, 2023) is displayed with a vertical red line. **B)** Temporal distribution of HPAI H5 (clade 2.3.4.4b) poultry farm outbreaks in four regions: region 1 (red): Norway (NW), Sweden (SW), Finland (FN); region 2 (green): Germany (GE), Denmark (DK), The Netherlands (NL), Belgium (BE); region 3 (yellow): United Kingdom (UK), Ireland (IR); region 4 (blue): Bulgaria (BG), Romania (RO), Hungary (HU), Poland (PO), Czech Republic (CZ). **C)** Temporal distribution of HPAI H5 (clade 2.3.4.4b) wild bird cases in four regions: region 1 (red): Norway (NW), Sweden (SW), Finland (FN); region 2 (green): Germany (GE), Denmark (DK), The Netherlands (NL), Belgium (BE); region 3 (yellow): United Kingdom (UK), Ireland (IR); region 4 (blue): Bulgaria (BG), Romania (RO), Hungary (HU), Poland (PO), Czech Republic (CZ). **D)** Predicted number of HPAI H5 poultry farm outbreaks in France as a function of the predictor variable: number of HPAI H5 wild bird cases in region 1: Norway (NW), Sweden (SW), Finland (FN). Black dots represent the observed number of French outbreaks in 2016-23. **E)** Predicted number of HPAI H5 poultry farm outbreaks in France and in heavily affected, non-vaccinating European countries in 2023-24. Horizontal black lines represent the corresponding 95% prediction intervals. Red stars represent the observed number of HPAI H5 poultry farm outbreaks in each country in 2023-24.

To explore this, we compared the observed reduction in outbreaks to what would have been expected based on historical outbreak patterns and external infection pressure, and considered potential factors that could explain any differences. We extracted surveillance data on HPAI H5 clade 2.3.4.4b detections in Europe (2016-24) from the FAO’s global animal health database (https://empres-i.apps.fao.org) (**Figure 1A-C**). For each epidemiological year (from September 1 to August 31), we retrieved poultry outbreak numbers in France.

Candidate predictors were defined using poultry outbreak and wild bird case numbers from neighboring countries as proxies for external infection pressure, based on evidence from phylogenetic showing genetic links between circulating viruses in these regions^5,6^. These predictors were defined for combinations of time windows (one, two or three months prior to the first reported poultry outbreak in France) and geographical regions (region 1: Norway, Sweden, Finland; region 2: Germany, Denmark, The Netherlands, Belgium; region 3: United Kingdom, Ireland; region 4: Bulgaria, Romania, Hungary, Poland, Czech Republic) (**Appendix, Table 1**). We assumed consistent surveillance efforts across years, which is likely valid for poultry due to standardized programs in Europe, but less certain for wild birds given the opportunistic nature of passive surveillance. Using quasi-Poisson univariate regressions, we identified the predictor most statistically associated with the yearly outbreak numbers reported in France during the pre-vaccination period 2016-23 (p-value < 0.05) with the strongest model fit (pseudo-R^2^ > 0.80). The 2023-24 value of that predictor was then used to predict the number of outbreaks that would have occurred in France in 2023-24, assuming no changes in mitigation strategies. To validate our statistical approach, we independently applied the same methodology to identify predictors best explaining outbreak numbers in heavily affected and non-vaccinating European countries in 2016-23 (**Appendix, Table 2 and Figure 1**), and used them to predict the number of outbreaks in 2023-24 in each of these countries separately. Finally, we compared the 2023-24 predictions with the observed outbreak numbers for all countries.

The best predictor of the number of poultry outbreaks in France was the number of reported wild bird cases in region 1 one month prior to the first reported outbreak in France.

This association does not imply direct causation but likely reflects overall infection pressure and spillover risk. Using this variable, the model predicted a 95% confidence interval of 314-756 outbreaks in France in 2023-24 (**Figure 1D**), significantly higher than the observed 10 outbreaks, and corresponding to a relative reduction of 96-99%. By contrast, predictions for the other countries aligned well with the observed outbreak numbers, supporting the validity of the statistical approach (**Figure 1E**). Of note in Germany, the observed number of outbreaks was at the lower end of the predicted range. Potential explanations could include improved implementation of biosecurity measures or changes in poultry population dynamics, which remain to be investigated.

Our findings suggest that the observed reduction in outbreak numbers in France for 2023-24 resulted from a major intervention not present in other European countries. Among the potential explanations, a decreased infection pressure in Europe^1^ would have uniformly affected our predictions across all countries and cannot explain the discrepancy only observed for France. Farm biosecurity compliance in France, which has gradually improved since 2016, did not undergo major changes in 2023-24 to explain the observed reduction^7^. Other measures, such as movement restrictions^8^ and indoor confinement^9^, also remained largely unchanged. While the duck population declined in 2020-22 due to prior outbreaks, it increased again in 2023^10^, ruling out a reduced duck population as an explanation. In this context, the vaccination campaign remains the most likely contributor to the reduction in outbreak numbers in France in 2023-24. Uncertainties persist regarding the potential indirect protection of unvaccinated poultry as well as other contributing factors, such as the potential role of virus evolution in virulence, which require further investigation^10^. Also, further mechanistic modeling of vaccination coverage would help quantify the direct impact of vaccination on outbreaks in more detail. Notably, vaccination should be implemented as part of a comprehensive approach that integrates biosecurity measures, regulation of poultry densities, and sustained virus surveillance, given the constant and unpredictable evolution of HPAI viruses.

## Statements

## Ethical statement

No ethical approval was needed for the study.

## Funding statement

This study was performed in the framework of the Chair for Avian Health and Biosecurity, hosted by the National Veterinary College of Toulouse and funded by the French Ministry of Agriculture and Food, General Directorate for Food.

## Use of artificial intelligence tools

None declared.

## Data availability

Data used in this study is available from the FAO’s global animal health database (https://empres-i.apps.fao.org)

## Preprint

https://www.biorxiv.org/content/10.1101/2024.08.28.609837v2

## Acknowledgements

No acknowledgements.

## Conflict of interest

None declared.

## Authors’ contributions

Study design: C.G., L.F., T.V. Data resources: L.F. Data analysis: C.G., L.F., E.M. Analysis guidance: S.L., T.V. Manuscript preparation: C.G. Review and approval of final manuscript: all authors

## Notes

### Competing Interest Statement

The authors have declared no competing interest.

### Summary of Updates

Figure 1 has been revised, text has been rephrased to clarify the methods

